# Environmental heterogeneity ruling the action of ecological processes over the bacterial metacommunity assembly

**DOI:** 10.1101/2020.01.21.914234

**Authors:** Paula Huber, Sebastian Metz, Fernando Unrein, Gisela Mayora, Hugo Sarmento, Melina Devercelli

## Abstract

Determining which processes take place in the spatial distribution of bacterioplankton metacommunities has been a central goal of microbial ecology. In freshwater ecosystems, selection has been spotted as the main driver shaping bacterial community. However, its relative importance compared with others processes (dispersal, drift, diversification) may depend on spatial heterogeneity and dispersal rates within a metacommunity. Still, a decrease in the role of selection is expected with increasing dispersal homogenization. Here, we investigate the main ecological processes modulating the bacterial assembly in contrasting scenarios of environmental heterogeneity. We carried out a spatiotemporal survey in the floodplain system of Paraná River. The bacterioplankton metacommunity was studied by a step-forward analysis combining different statistical inferences based on the phylogenetic and taxa turnover as well as co-occurrence networks. We found that selection was the main process even at both extremes of environmental heterogeneity and homogeneity, challenging the general view that the strength of selection is weakened due to dispersal homogenization. The ecological processes acting on the community also determined the complexity and randomness of bacterial networks. The heterogeneous selection promoted greater network complexity increasing the β-diversity, and bacterial associations were more random with the increasing influence of stochasticity. Finally, the spatiotemporal heterogeneity was an important factor determining the number and identity of keystone taxa in the system. Integrating all these empirical evidences we propose a new conceptual model that elucidates how the environmental heterogeneity guides the action of ecological processes shaping the bacterial metacommunity.

## INTRODUCTION

Determining the processes behind community’s assembly across Earth’s ecosystems is a major research topic in ecology [1–4]. Two perspectives address this issue. On one hand, the traditional niche-based theory states that deterministic processes such as environmental filters and species interactions, govern community structure and determine species composition [5, 6]. On the other hand, the neutral theory considers that all species are ecologically equivalent and, therefore, the structure of communities is a consequence of stochastic processes of birth, death, colonization, immigration, speciation, and probabilistic dispersal [1, 7, 8]. Although these perspectives are often considered contradictory, they are not mutually exclusive and a central debate is on their relative importance in shaping natural communities.

The metacommunity concept provides an adequate framework to identify these processes considering multiple spatial scales [9, 10]. Under this approach, communities are spatially connected in a network by dispersal, and the processes involved in their assembly are the result of an interplay between local factors and regional dynamics.

More recently, Vellend [3, 11] proposed a conceptual framework in which the structure of communities could be explained by four high-level processes: selection, dispersal, ecological drift, and diversification. Briefly, selection involves deterministic factors that modify the community structure due to fitness differences among individuals. It can act under homogeneous or heterogeneous environmental conditions, leading to more similar or dissimilar structure among communities (homogeneous or heterogeneous selection) [12, 13]. Dispersal results from both deterministic and stochastic factors that either favor or limit the movement (active or passive) and establishment of organisms among communities. High dispersal rates have a homogenizing effect (homogeneous dispersal) leading to a low turnover within communities, whereas low dispersal rates (dispersal limitation), coupled with drift or weak selection, increase community turnover [12, 14]. Drift influences changes in communities due to demographic events of birth, death and reproduction that occur at random, independently of species rates [15]. Finally, diversification refers to the generation of new species by genetic variation mainly due to stochastic factors, and a larger spatial and temporal scale with respect to the other processes are needed to observe its effects [16].

Bacteria are the most conspicuous and diverse components of the ecosystems’ microbiome, and the study of the processes shaping its structure contributes to gain proper comprehension of the functional organization of ecosystems. The ecological rules governing their metacommunity assembly change from land to water [17–21]. Differences among ecosystems are fundamentally due to environmental heterogeneity, connectivity, and spatial extent. Within aquatic ecosystems, the complexity of freshwater bodies is comparatively high comparing to oceans [22], consequently, striking idiosyncratic patterns in bacterial diversity were observed [23–27].

In freshwater ecosystems, deterministic processes (mainly selection) have been spotted as the main drivers shaping bacterial community structure [18, 26, 28–30]. However, it is not a general rule and the relative importance of selection compared with other processes may depend on the environmental heterogeneity and dispersion rates within a metacommunity [23, 27, 28, 31, 32]. For example, in systems with low environmental heterogeneity the stochastic processes seem to be more relevant, because environmental filters are not strong enough to act as selective forces allowing species sorting [31, 33]. In addition, bacterial communities in isolated systems should be mainly structured by dispersal limitation, whereas as environmental connectivity increases, homogeneous dispersal should become more important [29, 30, 33, 34].

Most previous results have been based on a metacommunity snapshot. However, in complex network systems subject to hydrological influence, the relative importance of the structuring ecological processes could change over time according to the variability in connectivity [29, 35–40]. Moreover, very few studies actually applied Vellend’s approach to its full extent [12, 30, 41– 43]. Another gap lies in understanding how the ecological processes influence the interactions between taxa (e.g. co-occurrence networks). Changes in community composition should affect the microbial co-occurrence networks [21, 44–49], thus we might expect more random co-occurrence and less complex networks when stochastic processes predominate. However, the empirical evidence of this dependency remains scarce.

The Paraná River floodplain constitutes a network of environments that resembles metacommunity organization [37], being an ideal system to address these issues. It is characterized by a wide range of temporal and spatial heterogeneity mediated by irregular hydrological fluctuations [50]. It comprises multiple shallow lakes, some of which are permanently connected to the main river or secondary channels, whereas others remain isolated still high water phases connects most the environments [51]. The floods have a homogenization effect on the environmental features [52, 53], as observed for other large rivers [54]. Here, we investigated the bacterial community structure through 16S amplicon sequencing in this complex floodplain system in order to (i) determine the relative importance of the High-level processes structuring the bacterial metacommunity, (ii) detect changes in bacterial co-occurrence networks, and (iii) identify keystone taxa in contrasting environmental heterogeneity scenarios mediated by landscape connectivity.

We hypothesized that the influence of different ecological processes shaping bacterioplankton metacommunities depend mainly on system’s environmental heterogeneity determined by the degree of landscape connectivity. To test this hypothesis, we first analyzed the processes shaping the community structure of bacterioplankton using the approach based on phylogenetic and taxa turnover proposed by Stegen et al. [12]. Then, we constructed co-occurrence networks to explore how ecological processes affect putative interactions between taxa. We predicted that, as the degree of connectivity increases and the system becomes more homogeneous, non-selective processes would have major relative importance in structuring the bacterioplankton metacommunity leading to more random associations between bacteria taxa.

## Material and Methods

### Study site and sampling design

The Paraná River is the second largest river in South America and fifth in the world [51] (Fig. 1). The middle stretch is formed by a main channel and a large floodplain that encompasses a high number of temporary and permanent streams and lakes. The system is characterized by a complex spatiotemporal dynamic influenced by hydrosedimentological regime. It has a fluctuating hydrological dynamic with pulses of flood and drought that determine high and low water phases, and alternate in variable frequency with extreme hydrological phases [55]. During high water phases, water drains to the floodplain, connecting the environments in different degrees according to the magnitude of the flood. The sedimentary pulse that mainly depends on the suspended solids coming from the Bermejo Basin, is not coupled with the hydrological pulses, affecting abiotic factors of the environments connected with the main channel [56]. The dynamics of biological communities respond to a great extent to the hydrological fluctuations, since it has a direct influence on the environmental characteristics, dispersal processes, and habitats colonization [36–39, 57–59].

**Fig. 1:**
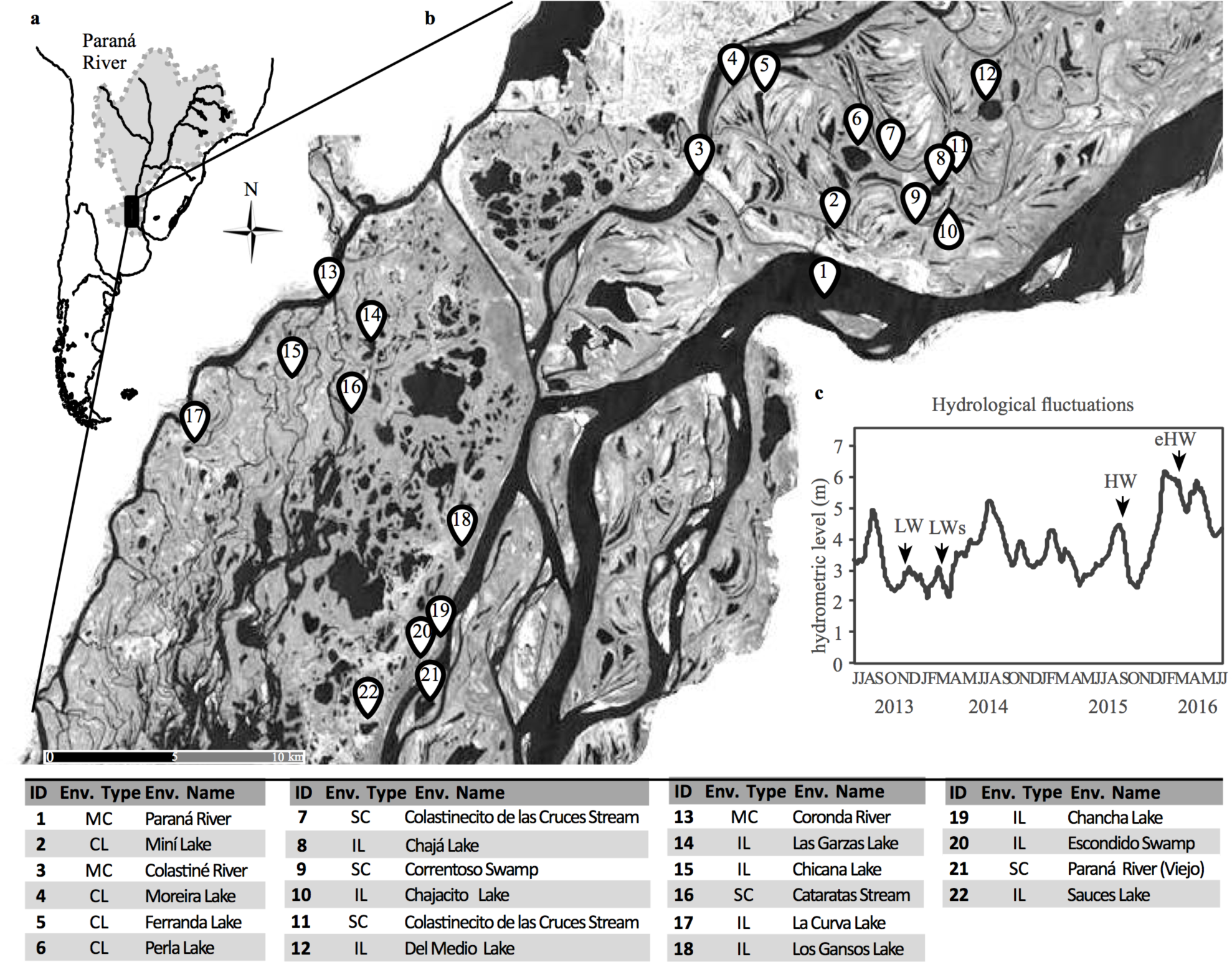
**a)** The Paraná River in South America showing the location of the study area and **b)** the environments sampled. **c)** Daily water level from 2013 to 2016 in the Paraná River. The sampled periods are indicated with arrows.

The temporal dynamic and the spatial heterogeneity of the system were captured sampling different hydrological phases and environment types characteristics of the floodplain River (Fig. 1). Four samplings that lasted 10 days each were performed during: a low-water phase (LW, November–December 2013), a low-water phase coupled with the sedimentological pulse (LWs, March–April 2014), a high water phase (HW, September 2105), and an extraordinary high water phase (eHW, March 2016). Four representative environment types with different hydrological connectivity and morphological features were selected (Fig. 1): the main channel and large secondary channels (MC), minor secondary channels (SC), connected lakes (CL), and isolated lakes and swamps (IL). Lentic environments were classified as IL or CL according to its hydrological connectivity degree during low water phase. Overall our dataset was constituted by 59 samples: 16 from LW, 17 from LWs, 13 from HW and 13 from eHW.

#### Environmental data collection

Depth, water temperature, pH, conductivity, dissolved oxygen (DO) concentration (HANNA checkers), and Secchi disk depth were measured in situ. Subsurface water samples were collected and transported in polypropylene containers to the laboratory for turbidity, soluble reactive phosphorus (SRP), nitrate (NO_3_^-^), ammonium (NH_4_^+^), chromophoric dissolved organic matter (CDOM) concentration (A_440_), CDOM molecular weight (S_275-295_), and chlorophyll-*a* analyses. Technical details of the abiotic variables’ analyses are described in the Supplementary Methods.

#### Bacterial samples collection and sequencing

Subsurface water samples for DNA (100 to 140 ml) were prefiltered with a 50 µm pore mesh, and filtered through a 0.22 µm pore-size polycarbonate filters (Millipore). The filters were frozen (−80°C) until DNA extraction. Genomic DNA was extracted using a CTAB protocol [60] as described in the Supplementary Methods.

Tagged amplicons of the 16S rRNA gene (V3–V4 region) were obtained with the primers 341F and 805R [61] and sequenced using Illumina MiSeq 2×250 paired-end reads approach [62] by Macrogen (Seoul, South Korea).

#### Sequence data analysis

Raw sequences were processed using a modified version of the pipeline proposed by Logares [63] (https://github.com/ramalok), described in the Supplementary Methods. Operational taxonomic units (OTUs) were defined with no clustering (zero-radius OTUs [zOTUs]) using UNOISE2 algorithm [64]. The zOTU table resulted in 10 253 zOTUs (2 560 053 reads). We constructed zOTU rarefaction curves to evaluate richness saturation (Fig. S2).

A second zOTU table was generated discarding zOTUs with <10 reads as well those assigned to Archaea domain or chloroplast. This table was normalized to an equal sampling depth to generate the subsampled zOTU table (rrarefy function, Vegan v.2.0.9 package [65] in the R environment [66]) which consisted of 8 927 OTUs and 18 747 reads per sample.

The sequence data obtained in this work were deposited at the European Nucleotide Archive (ENA) public database with the references ERSXXXXXXX to ERSXXXXXXX.

#### Environmental heterogeneity

The environmental heterogeneity was evaluated by computing the average dissimilarity between sites (Ed) [67] based on nine abiotic variables: DO, turbidity, conductivity, pH, SRP, NO_3_^-^, NH_4_^+^, CDOM concentration, and CDOM molecular weight. For each hydrological phase we computed a Euclidean distance matrix (Vegan package, R) and calculated the dissimilarity between sites (*Ed*) as follow:

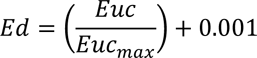

where *Euc* is the Euclidean distance between two sites and *Euc_max_* corresponds to the maximum Euclidean distance considering all the pairwise distances in the overall dataset. 0.001 was added to account for zero similarity between sites [67]. Then, we calculated the mean *Ed* (Ed) of each similarity matrix computed and used it as an index of environmental heterogeneity in each hydrological phase. Additionally, for each hydrological phase we calculated the coefficient of variation (CV%), as the quotient of the standard deviation and mean of each variable.

To test if the environmental conditions in the four hydrological phases differed significantly, we performed a permutational multivariate analysis of variance (PERMANOVA) [68], using Euclidean similarity index, and Bonferroni pairwise post-hoc tests (Past software V3 [69]).

#### Bacterial community structure and diversity

To a general view of bacterial community, we analyzed the taxonomic structure in each hydrological phase and constructed rank abundance curves (Fig. S3). We characterized the “rare biosphere” and the “abundant fraction” (zOTUs relative abundances per sample <1% and >1%, respectively) [70].

The effect of hydrological conditions on bacterial community structure was assessed by calculating the community turnover in each hydrological phase. We computed a Bray-Curtis distance matrix on the basis of zOTU normalized abundance for each hydrological phase and calculated the dissimilarity (*Xd*) between the local communities as follow:

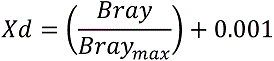

where *Bray* is the Bray-Curtis distance between two communities and *Bray_max_* corresponds to the maximum Bray-Curtis distance considering the overall dataset. Then, we calculated the mean *Xd* (Xd) of each similarity matrix computed, which was used as a value of community turnover.

We also evaluated the significance of the effect of hydrological conditions on bacterial turnover performing a PERMANOVA [68] with 9 999 permutations on Bray-Curtis dissimilarity distance, and Bonferroni pair-wise comparisons were run to obtain *P*-values (Past software V3).

To visualize the taxonomic similarity across local communities, nonmetric multidimensional scaling (NMDS) was performed using the Bray-Curtis distance metric (Vegan package, R). zOTU richness, Shannon-Weaver diversity (H’) and Whittaker β-diversity indices were calculated from the normalized bacteria zOTU table (Past software V3). To test whether both attributes differed significantly (*P*<0.01) among hydrological phases we performed a Kruskal-Wallis test with Mann-Whitney U test.

#### Quantification of high-level process structuring the bacterial metacommunity

To quantify the relative importance of ecological processes in structuring the bacterial metacommunity we used the approach proposed by Stegen et al. [12]. This method analyzes the influence of environmental filtering based on an intrinsic property of the community, independently of the considered environmental variables [12, 71], sorting the problem of overestimating the effect of stochastic processes due to unmeasured environmental variables.

We first measured the influence of selection in each hydrological phase, comparing observed phylogenetic turnover to a random expectation using the βNTI Index (β-Nearest Taxon [72–74]). This metric is defined as the difference between the observed mean phylogenetic distance between each taxon and its closest relative in two communities (βMNTD metric) with the βMNTD obtained from the null distribution, divided by the standard deviation of phylogenetic distances in the null data. Absolute βNTI value greater than 2 (|βNTI |>2) indicates coexisting taxa are more closely related than expected by chance, so selection strongly influences community composition [12, 75]. Next, we estimated the percentage of homogeneous selection as the fraction of pairwise comparisons with a βNTI value of <−2 and heterogeneous selection as the fraction of pairwise comparisons with a βNTI value of >+2 [12].

As this approach considered that the habitat preferences of closely related taxa are more similar than the habitat preferences of distantly related taxa [12], we tested the phylogenetic signal performing a Mantel correlogram analysis between zOTU niche and zOTU phylogenetic distances [12, 23, 41, 76–78] (see Supplementary Methods). Phylogenetic signals were detected over short phylogenetic distances (Supplementary Fig. S1) consistent with previous work [12, 23, 41, 78, 79].

The communities that were not structured by selection (i.e. |βNTI |<2), were then analyzed in the second step, where the action of dispersal and drift was calculated based on the taxonomic (zOTU) turnover with the Raup-Crick metric [80] using Bray-Curtis dissimilarities (hereafter RC_bray_) [12]. RC_bray_ compares the measured β-diversity from a metacommunity against the β-diversity obtained from a null model. RC_bray_ values between –0.95 and +0.95 indicate significant departures from the degree of turnover, occurring when drift is acting alone [12]. Additionally, RC_bray_ values >+0.95 indicate that communities are less similar than expected by chance as a result of dispersal limitation combined with drift, and RC_bray_ values <–0.95 indicate that communities are more similar than expected by chance as a result of homogeneous dispersion [12].

#### Co-occurrence networks analysis

To infer whether bacterial associations were influenced by the different ecological processes structuring the metacommunity, we performed a network analysis using the SparCC algorithm [81] implemented in FastSpar [82]. One specific network association for each hydrological phase was constructed considering zOTUs with >50 reads and present in at least 2 samples. FastSpar was run using 1 000 interactions and 1 000 bootstraps to calculate the *P*-values. Only the significant (*P*<0.01) and strong positive and negative correlations (| r | >0.7) were considered. The networks were visualized in Cytoscape V3.7.1 [83]. To validate the non-random co-occurrence patterns, we evaluated networks against their randomized versions using the Barabasi–Albert model available in Randomnet-works plugin in Cytoscape. NetworkAnalyzer tool [84] was used to calculate five network topology properties: number of nodes (bacterial zOTUs), number of edges (associations between bacteria), average number of neighbors, distribution of degree and clustering coefficient. These structural attributes were used to infer the complexity and chance in bacterial associations [46, 85–87]. The average number of neighbors is the average connectivity of a node in the network, thus higher values are expected according increase the complexity of the associations [88]. The distribution of degree indicates the number of nodes that have each degree, so the complexity of the associations can be inferred by analyzing the power-law distribution [46, 88]. Finally, the clustering coefficient indicates how nodes are embedded in their neighborhood and, thus, the degree to which they tend to cluster together [88, 89]. As consequence, low values of this metric are interpreted as more chance in the association patterns.

#### Keystone bacterial taxa

We studied those taxa that play central roles on community functioning, commonly referred as keystone species [90, 91]. For each network we identified the keystone taxa as those nodes with a high degree (>30), closeness centrality (>0.36) and low betweenness centrality (<0.07) [46, 90, 92] (NetworkAnalyzer tool in Cytoscape). These metrics illustrate both the number of connections and how important those connections are to the overall network [90, 93, 94].

Redundancy Analysis (RDA) was performed to evaluate the effect of abiotic variables on the abundance of the keystone taxa (CANOCO software V5 [95]). Environmental variables tested as explanatory variables (DO, turbidity, conductivity, pH, SRP, NO_3_^-^, NH_4_^+^, CDOM concentration CDOM molecular weight), were normalized as standard normal deviates. A forward selection procedure was run to find the subset of the significant explanatory variables using the Monte Carlo Permutation.

## RESULTS

### Environmental heterogeneity

The average environmental dissimilarity between sites (Ed) as well coefficient of variation (CV) of the nine abiotic variables indicated a clear trend to spatial homogenization from low water (low hydrological connectivity) to high water (high hydrological connectivity) phases (Fig. 2). Additionally, the four hydrological phases were significantly different according to their environmental characteristics (PERMANOVA *P*<0.05, Table S1 and S2).

**Fig. 2:**
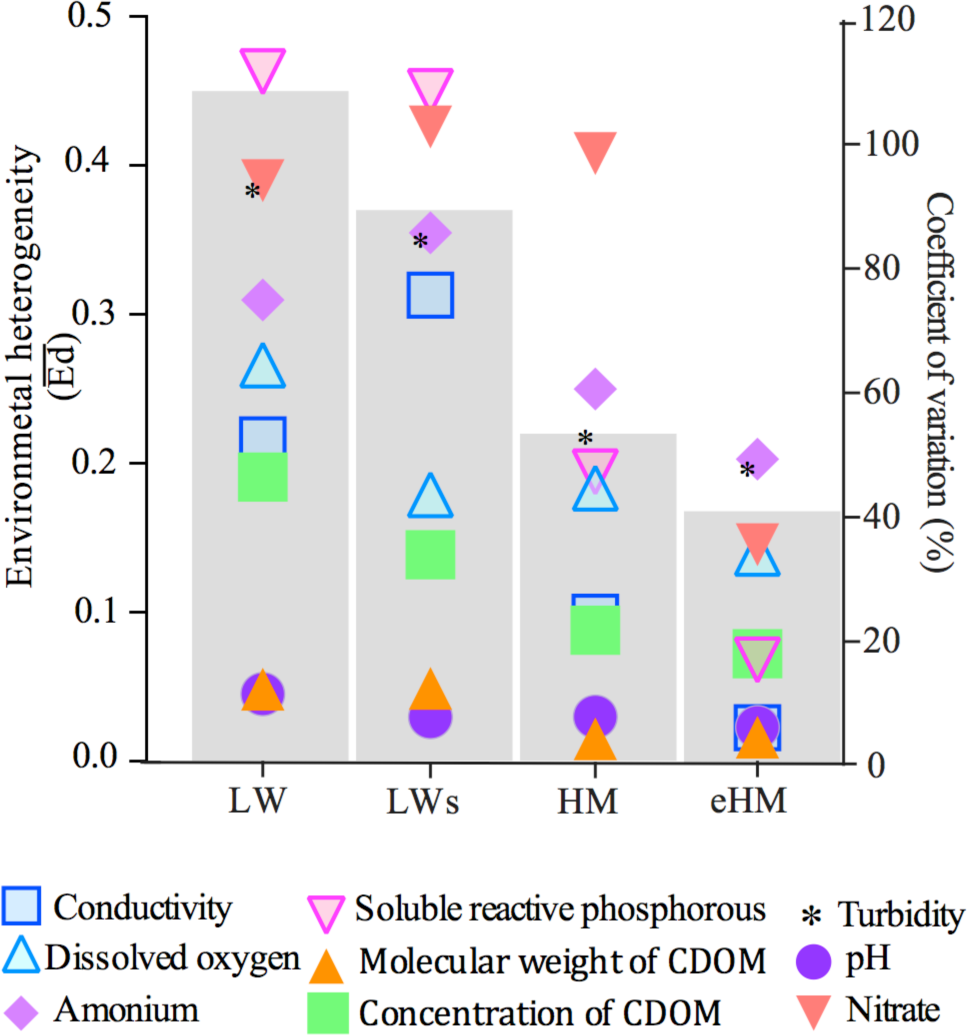
Environmental heterogeneity in the Paraná fluvial system at each hydrological phase expressed as average environmental dissimilarity between sites (Ed, grey bars) and the variability (coefficient of variation, CV%) of nine selected abiotic variables among environments. **LW:** low water, **LWs:** low water with sedimentological pulse, **HW:** high water, **eHW:** extraordinary high water.

### Bacterial community structure and diversity

Overall in the 59 samples, a total of 8 927 zOTUs within the bacterial domain were defined. The metacommunity was mainly dominated by Proteobacteria (33%), Actinobacteria (23%), Bacteroidetes (18%), and Verrucomicrobia (11%). The community structure according to zOTUs abundance varied with hydrological conditions with significant differences among phases (PERMANOVA, *P*<0.001, Fig. S4, Table S3) except between low water phases (LW and LWs). In the NMDS, samples were organized following a clear homogenization pattern from low water to high water phases (Fig. 3). In the eHW phase, local communities were more similar displaying a distinct cluster. In contrast, those from low water phases were more spread, mainly due to differences in the community composition of isolated lakes (Fig. 3). The community turnover (Xd) varied markedly among hydrological phases, being significantly lower (PERMANOVA, *P*<0.005) in high water phases (Xd: HW=0.58; eHW=0.29) than in low water phases (Xd: LW=0.76; LWs=0.75). In line with this, the highest values of ß-diversity were obtained in both low water phases (Whittaker: LW=3.50; LWs=3.27; HW=1.97; eHW=1.95).

**Fig. 3:**
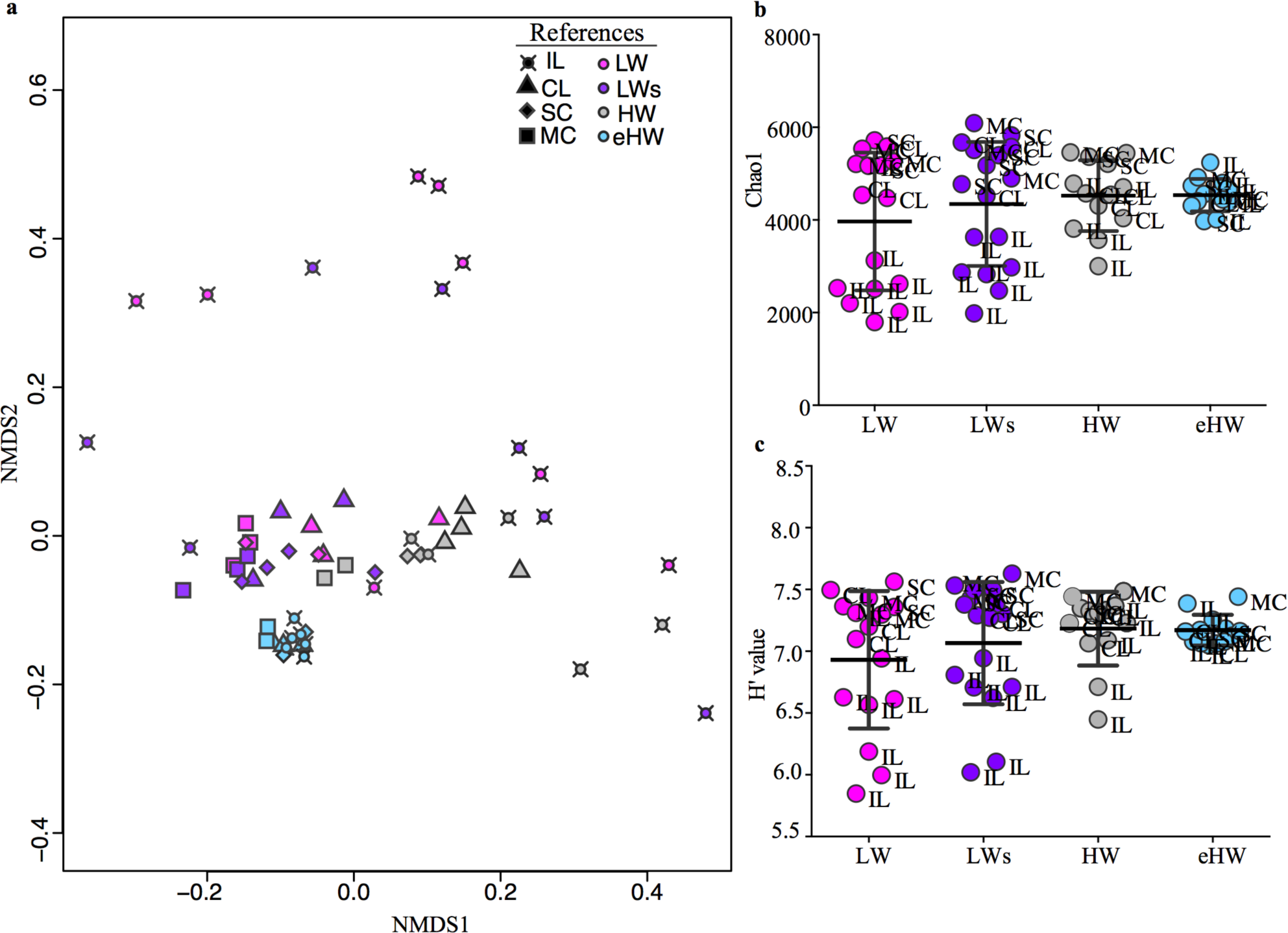
a) Samples ordination in a non-metric multidimensional scaling (NMDS) according to the similarity in bacterial communities structure (zOTU relative abundance) among four hydrological phases (indicated with color) at the different type of environments (indicated with symbols) of the Paraná fluvial system. Stress value: 0.112. **b-c)** Richness and Shannon-Weaver diversity index H’ of local bacterial communities at the Paraná fluvial system in four hydrological phases. **LW:** low water, **LWs:** low water with sedimentological pulse, **HW:** high water, **eHW:** extraordinary high water. **IL:** isolated lake, **CL:** connected lake, **SC**: secondary channel, **MC:** main channel.

Species richness and Shannon diversity (H’) presented similar average values at the different hydrological phases (ANOVA, *P*=0.366, Fig. 3). However, differences among local communities were pronounced at low water phases, when isolated lakes were more spread showing lower values than the mean (Fig. 3).

### High-level process structuring the bacterial metacommunity

Having demonstrated that bacteria community was clearly linked to changes in the environmental heterogeneity, we quantified the relative importance of selection, dispersal, and drift processes in structuring the metacommunity.

The phylogenetic turnover analysis (βNTI) revealed that selection was the most important structuring process regardless hydrological periods. However, its relative importance as well as the type of selection, changed according to hydrological conditions and hence, to the environmental heterogeneity. During both low water phases, heterogeneous selection had the major role in structuring the metacommunity (76.66% LW and 84.55% LWs of the overall community turnover) being poorly significant the role of homogeneous selection and non-selection process. In HW phase, heterogeneous selection was also the most important process (39.74%), but the relative importance of homogenous selection and non-selection processes were twice as high compared to the low water phases (Fig. 4). In contrast, homogenous selection was the most important process in eHW (88.46%) (Fig. 4). Regarding non-selection processes (RC_bray_), dispersal limitation combined with drift had the prevalent role in all the hydrological phases, except in eHW period when homogeneous dispersion became more important (Fig. 4).

**Fig. 4:**
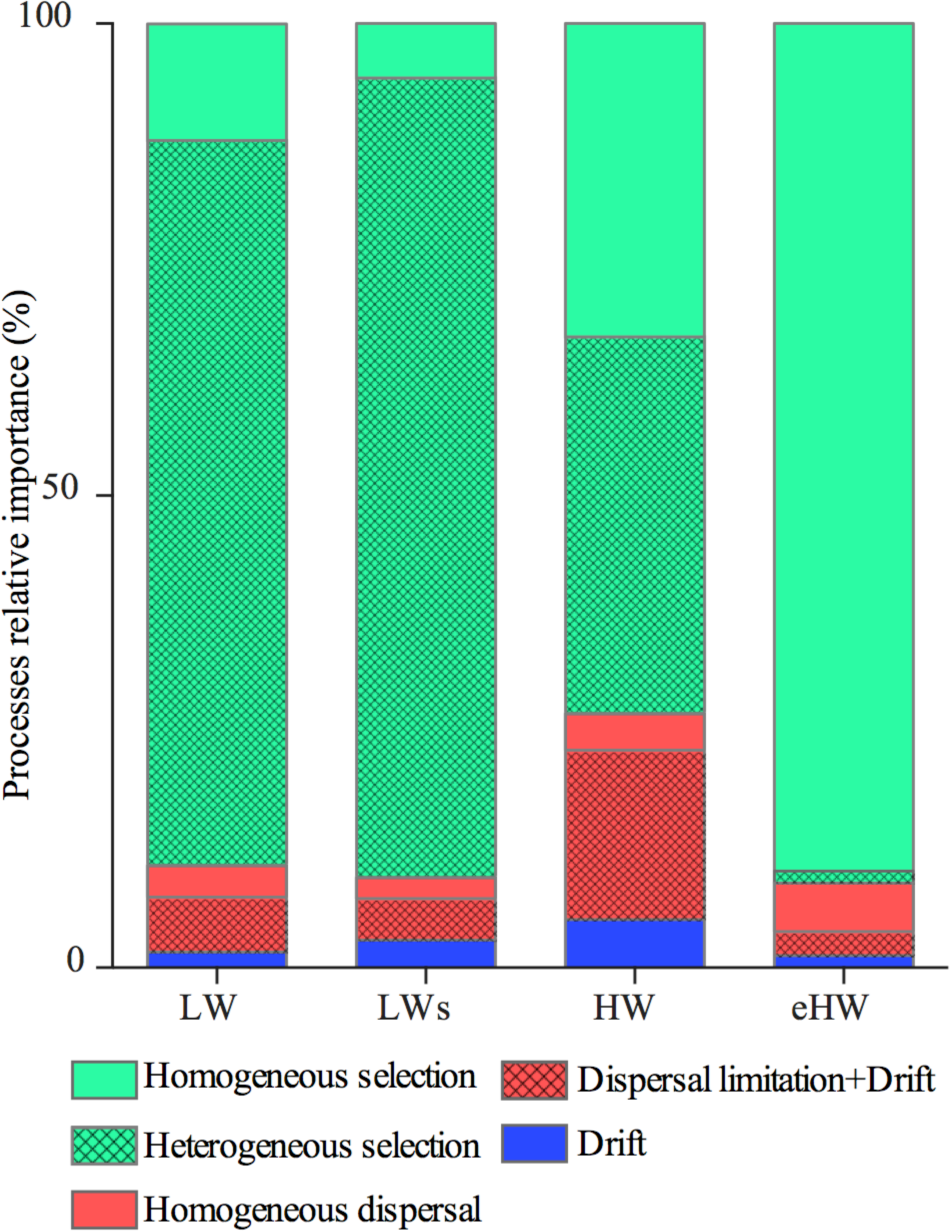
Relative importance of high-level processes structuring the bacterial metacommunity of the Paraná fluvial system in four hydrological phases. Values indicate percentage of community turnover associated to each process: homogeneous selection, heterogeneous selection, homogeneous dispersal, dispersal limitation combined with drift, and drift. **LW:** low water, **LWs:** low water with sedimentological pulse, **HW:** high water, **eHW:** extraordinary high water.

### Bacterial community associations

The associations between bacterial zOTUs remarkably differed regarding the environmental heterogeneity (Fig. 5). The networks varied in both, number of nodes (bacterial zOTUs) and edges (associations between bacteria), being the highest under the dominance of heterogeneous selection (low water phases) (Fig. 5). The higher network complexity was observed during the dominance of heterogeneous selection, as the average number of neighbors was considerably higher in both low water phases with respect to high water phases, especially HW (Fig. 5). In agreement, the histograms of degree distribution showed a stronger power-law in low than in high water phases (Fig. 6), indicating the presence of nodes highly connected when the heterogeneous selection had the major role in structuring the metacommunity.

**Fig. 5:**
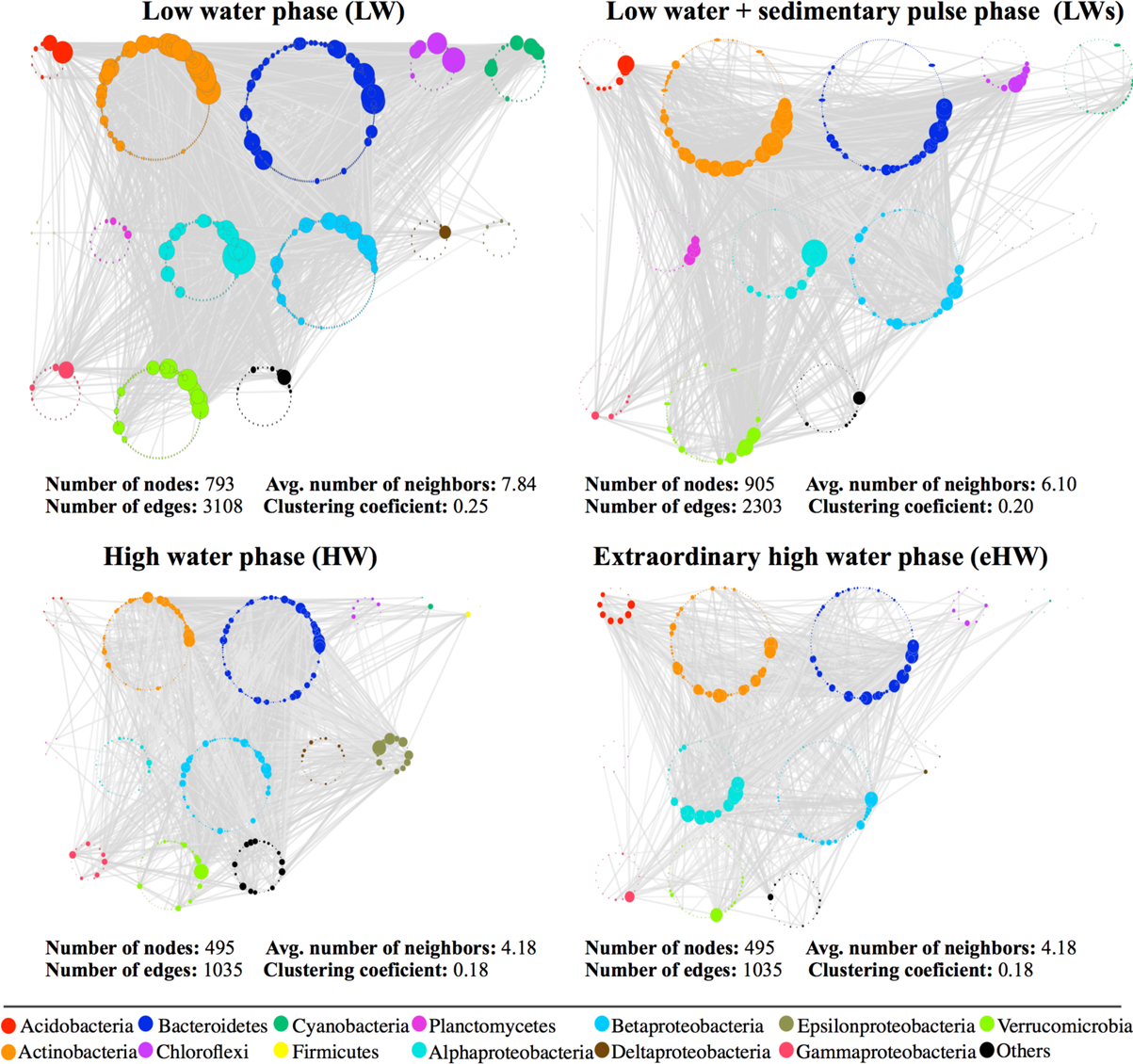
Bacterial associations networks for each hydrological phase of the Paraná fluvial system, arranged according to bacterial taxonomy (classes or orders). The size of each node is proportional to the number of connections (degree). Only significant (*P*<0.01) and strong correlations (| r | > 0.7) were considered. The number of nodes, number of edges, average number of neighbors, and clustering coefficient of the overall network was indicated below each network.

**Fig. 6:**
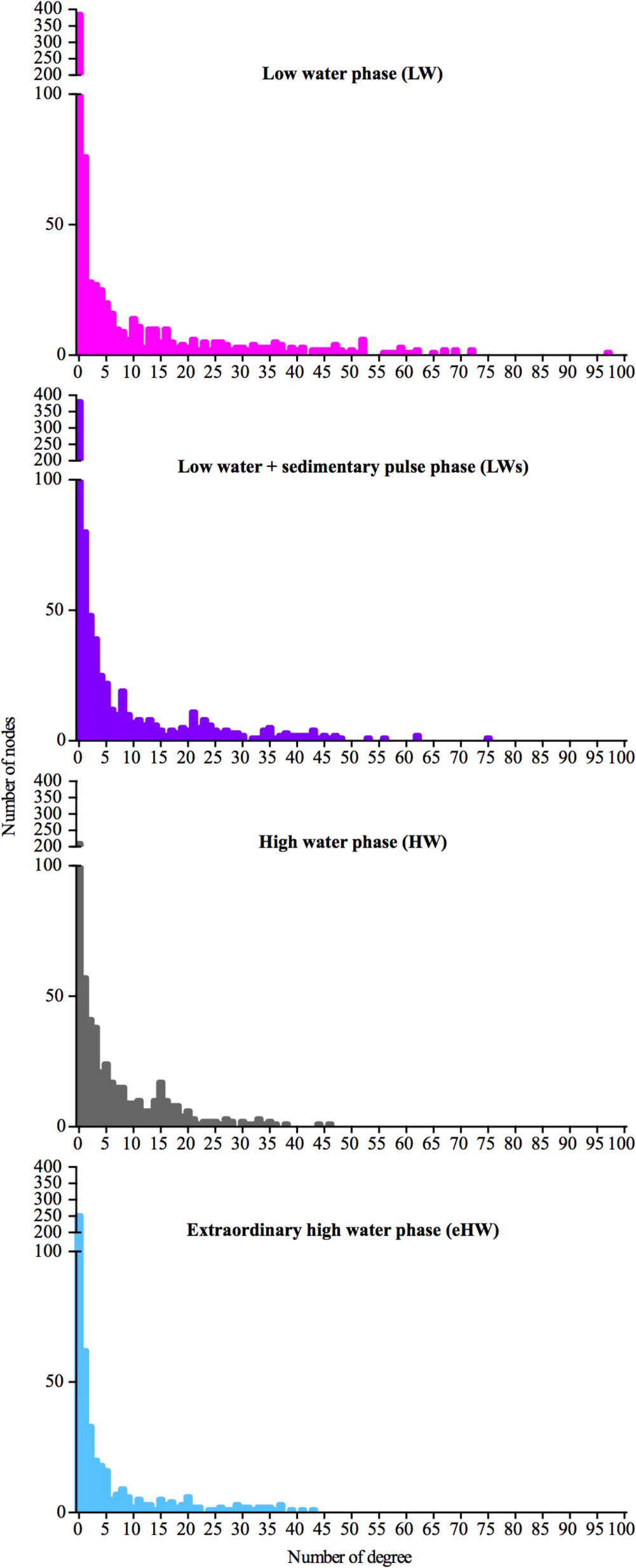
Complexity of bacterial associations networks for each hydrological phase of the Paraná fluvial system expressed as the distribution of degrees. Degree indicates the number of associations shared by each node in a network.

Finally, the values of clustering coefficient revealed more random associations during the HW phase (Fig. 5) when heterogeneous and homogeneous selection had similar importance in structuring the metacommunity, and the non-selection processes had higher contribution compared to the other phases.

### Metacommunity keystone taxa

The number as well as the taxonomic composition of keystone taxa were notably different in the four hydrological phases. In LW, 50 zOTUs were defined as keystones that were reduced to 11 in LWs and HW, and 13 in eHW. Most of these keystone taxa were exclusive of each hydrological phase and no one was present at the four phases. Keystone taxa accounted differential contributions to each phylum in each hydrological phase, being Actinobacteria and Bacteroidetes the better represented in all phases (Fig. S5).

In the RDA, keystone taxa were arranged according to the hydrological phase in which they were defined (Fig. 7). The first two axes accounted for 43.04% of the variance (axis 1: 33.09%; axis 2: 9.95%) being CDOM concentration (A_440_), CDOM molecular weight (S_275-295_), DO, and nutrients (SRP, NH_4_^+^ and NO_3_^-^) the main explanatory variables (*P*<0.01). The first axis was mainly defined by the NO_3_^-^ concentration and CDOM molecular weight (intra-set correlation coefficients: −0.75 and 0.65, respectively). The second axis was principally associated with CDOM concentration and molecular weight (intra-set correlation coefficients: 0.70 and −0.70, respectively), separating low from high water samples. The keystone taxa from both low water phases were positioned towards the upper part of the graph (Fig. 7). Taxa from the LWs grouped together by the left of the graph, while those from LW showed a clear segregation in 2 groups: taxa with higher abundances in lotic systems were positively related to NO_3_^-^ and NH_4_^+^ by the left panel; whereas taxa with higher abundance in lentic environments were inversely related with CDOM molecular weight by the right panel. The keystone taxa from high water phases were mainly positioned towards the lower part of the graph (Fig. 7): taxa from eHW located in the left panel were mainly associated with high CDOM molecular weight, meanwhile those from the HW positioned throughout the right panel were more related with high CDOM concentration (A_440_).

**Fig. 7:**
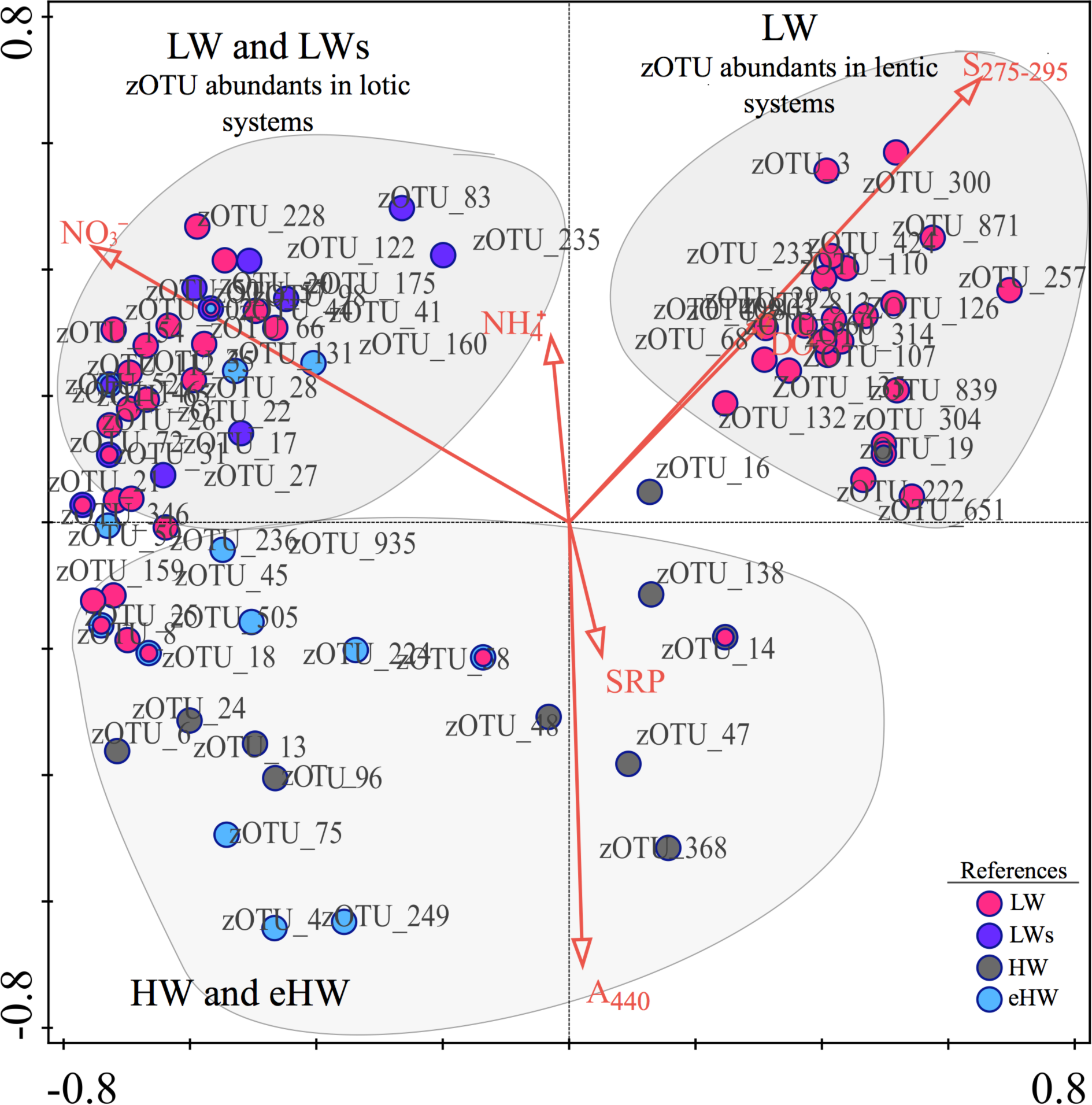
Redundancy analysis based on the abundance of keystone taxa per site versus environmental variables. Only the significant variables (*P*<0.05, forward selection) are shown (red arrows). CDOM (A_440_) is the light absorption coefficients at 440 nm that is proportional to the CDOM concentration. CDOM (S_275-295_) is spectral slope from 275 to 295 nm and indicates the CDOM molecular weight. Circles represent the position of each zOTU in the biplot and the color indicates the hydrological phase in which they were selected as keystone: low water (LW), low water with sedimentological pulse (LWs), high water (HW), extraordinary high water (eHW). Those zOTUs selected in more than one hydrological phase are indicated with different circles.

**Fig. 8:**
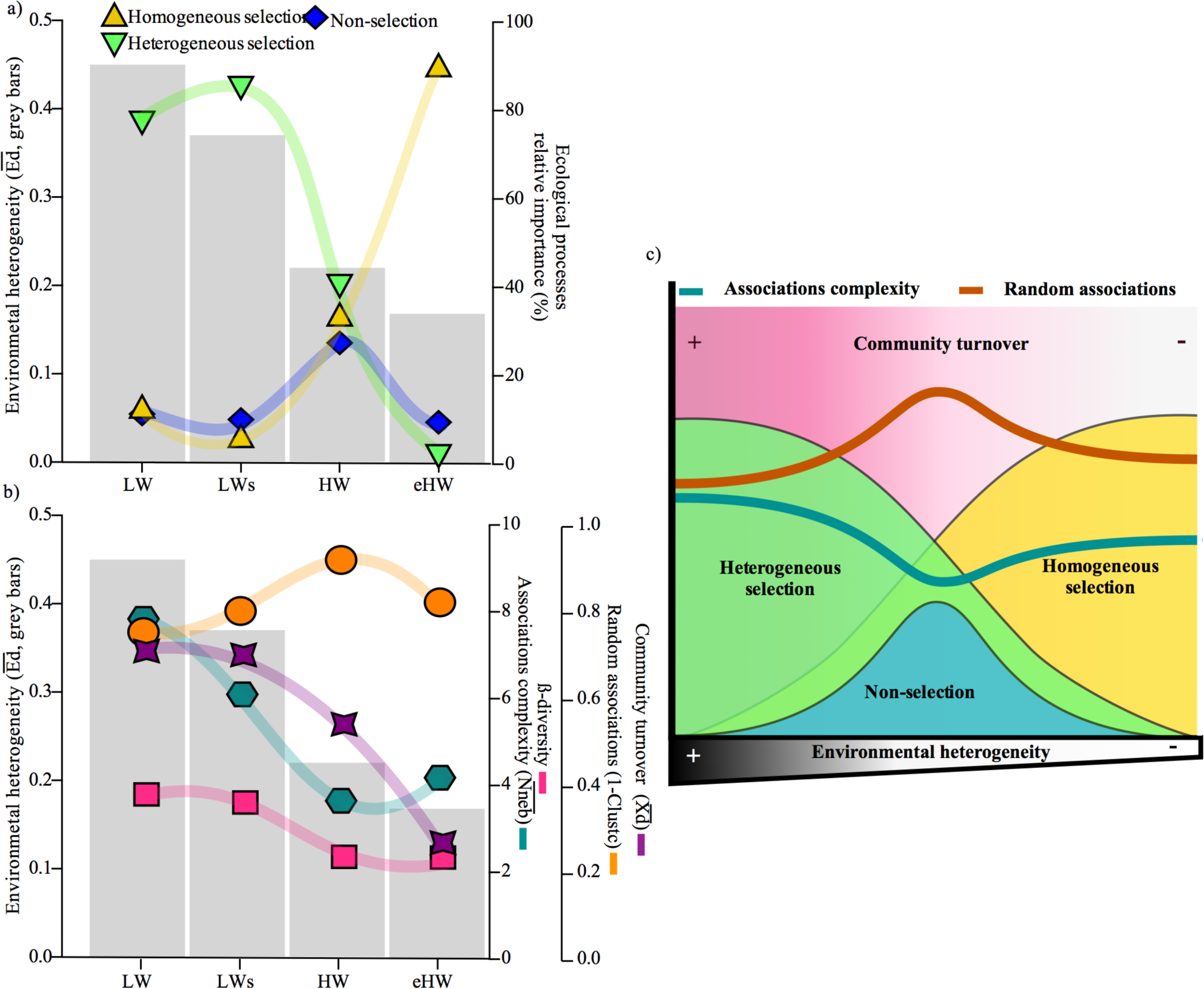
Empirical evidence **(a-b)** that support a new conceptual model **(c)** on how environmental heterogeneity regulates the four ecological processes that shape bacterial metacommunity assemblage (*sensu* Vellend [3, 11]). At high environmental heterogeneity, community structure is mainly determinate by heterogeneous selection. The heterogeneous selection promotes a greater β-diversity increasing divergence in local communities, and a greater complexity bacterial association networks. At intermediate values of environmental heterogeneity both heterogeneous and homogeneous selections act with similar strength, and stochastic processes reaches more importance. In this scenario, the selection acting in opposite ways does not allow homogenization or differentiation of local community structure, leading to an overall reduction of turnover and β-diversity. Simultaneously, the metacommunity would have a relatively high randomness due to intermediate dispersal rates. As consequence, the association network would tend to be less complex and more random. At extremely homogenous environmental conditions, the low diversity of niche habitat leads to an increase of homogeneous selection, and local communities are strongly filtered by common environmental factors. The homogeneous selection leads to a low turnover and β-diversity, promoting less complex associations.

## Discussion

This study provides the first empirical evidence that selection is the most important process structuring the bacterial metacommunity at both extremes of environmental heterogeneity and homogeneity. This finding challenges the general view that the strength of the selection is weakened due to dispersal homogenization. Additionally, at intermediary environmental heterogeneity, stochastic processes became more important leading to more random metacommunity network.

Based on the underpinning of Vellend’s approach [3], we predicted that the influence of non-selective processes would increase as the system becomes more homogeneous. Contrarily, we found that selection was the main process shaping the bacterial metacommunity not only at environmental heterogeneity but also at environmental homogeneity.

Several studies found evidence of the pivotal role of selection determining bacterial communities’ structure in different environments, and a decrease in its relative importance as dispersal rates increase and the systems become more homogeneous in their environmental conditions [29, 30, 78, 96]. Particularly, Wang et al. [23] in a comparative survey of a broad range of ecosystems (i.e. soil, stream biofilm, and lake), observed a clearly dominant role of deterministic processes on bacterial assemblages, whereas stochastic processes became more relevant as environmental heterogeneity decreased and dispersal rates increased. The authors proposed a theoretical threshold of selective strength above which communities should be mainly structured by deterministic processes as a consequence of high environmental heterogeneity and low dispersal rates.

However, these previous attempts only considering spatial scales. To address the full scope of the question, we extended these findings considering the spatial and temporal variability in a complex ecosystem. We summarized our findings on three possible scenarios: 1) With high environmental heterogeneity characterized by low connectivity, the structure of bacterial assemblage was mainly driven by heterogeneous selection (Fig. 8a). In this scenario, species with different fitness may be strongly filtered by different selective forces in each local community leading to a high β-diversity, whereas the influence of drift and dispersion should be irrelevant; 2) At intermediate environmental heterogeneity, both heterogeneous and homogeneous selections had similar weight and the relative influence of stochastic processes increased (Fig. 8a). Here, the magnitude of ordinary floods is insufficient to thoroughly mix water among all the floodplain, and some environments begin to homogenize in their environmental filters while others retain their identity [97]. The selection may act with similar strength both homogenizing and segregating local communities, leading to an overall reduction of community turnover and β-diversity. Simultaneously, the metacommunity should be relatively random due to intermediate dispersal rates; 3) At extremely environmental homogeneity, the homogeneous selection was the main ecological rule structuring the metacommunity (Fig. 8a). Despite stochasticity is expected to increase due to the high dispersal rates, the bacterial species should be strongly filtered by common environmental factors and the strength of homogeneous selection may dampen the influence of drift and dispersion. For instance, low community turnover and β-diversity are expected. These evidences give support to the early assumptions made by Stegen et al. [41] and Dini-Andreote et al. [78] and to a new view of bacterial community assembly in which the action of selective and non-selective processes are considered.

The understanding of ecological processes acting on metacommunities was extended studying the bacterial co-occurrence patterns. Network analyses-based approaches have the potential to infer inter-taxa correlations and can be applied to investigate structure complexity [47, 98], randomness [99], and the ecological rules guiding community assembly [44]. Particularly, the influence of ecological processes on bacterial interactions is a poorly explored field. Emerging studies in soil communities have revealed that systems under organic farming harbor more complex networks than conventional farming, mainly associated to higher habitat heterogeneity [46]. Additionally, it has been recently demonstrated that greater microbial diversity ensures greater association complexity [47].

Our results demonstrate that networks’ topology was markedly different according to the weight of the ecological processes acting on the bacterial community assemblage. At heterogeneous selection dominance (LW and LWs phases), the network associations had higher complexity (Fig. 8b). We assign this finding to the fact that the heterogeneous selection promotes a greater taxonomic β-diversity [47]. Contrarily, the homogeneous selection drove less complex networks decreasing the β-diversity. Additionally, under multiple ecological processes acting with similar strength, the network also tended to be less complex (Fig. 8b).

Another important finding of our study was the influence of the ecological processes on the randomness of networks. We observed less chance of bacterial associations when selection was stronger (Fig 8b). This is a general pattern mainly associated to the action of selection as taxa with similar ecological requirements tend to co-occur more than by chance (e.g. [44, 99, 100]). However, this is one of the first studies that have empirically linked bacterioplankton networks to structuring processes. Thus, our results add a new dimension revealing that both heterogeneous and homogeneous selection promotes less random associations. Additionally, as stochastic processes become more relevant, more random networks would be expected.

A useful feature of network analysis is that it allows to identify the strongly interconnected keystone taxa which have a core effect on the community assemblage [87, 90] and are optimal predictors of overall community changes [91]. Each hydrological phase presented a particular set of keystone taxa that were determined by different selective factors. It was not surprisingly considering that environmental conditions significantly varied among the hydrological phases (Fig. 1). Perhaps the most intriguing fact of our findings was the huge number of keystone taxa found under the highest environmental heterogeneity (50 vs 11 to 15 taxa). Even under similar networks’ complexity and structuring processes at both low water phases, the number of keystone taxa was clearly higher. We attribute this finding to a higher niche segregation and number of significant environmental factors related with keystone taxa from LW. While the inorganic dissolved nitrogen was the main driver at both low water periods, in LW the molecular weight of CDOM had also a significant role in determining keystone taxa, particularly those abundant in isolated lakes. This result brings new evidence to previously assumption about the role of environmental heterogeneity in determining the number and identity of keystone taxa [46, 90].

## Concluding remarks

Integrating all the empirical evidence obtained here we propose a conceptual model that synthetizes how environmental heterogeneity guides the action of ecological processes assembling the bacterial metacommunity (Fig 8c). In systems with high environmental heterogeneity, the heterogeneous selection plays the major role in structuring the community, promoting a greater β-diversity, and more complex association networks. At intermediate values of environmental heterogeneity, the action of stochastic processes reaches more importance, and both heterogeneous and homogeneous selections have similar contribution; this leads to a decrease of β-diversity and to more randomness in networks associations. In systems with high environmental homogeneity, the strength of homogeneous selection dampens the influence of drift and dispersion, and local communities become more similar and the association networks less complex.

The strength of this model is that it was conceived relying on different metacommunity features (e.g. taxonomic and phylogenetic turnover, network associations) taking into account the spatial and temporal scales. Furthermore, we used a step-forward analysis combining different statistical inferences, which allowed us to take advantage of the strengths of each method alone and in synergy with the others. Therefore, the proposed model represents a significant improvement on the fundamental knowledge of bacterial community’s assembly across freshwater ecosystems and provides a new framework for future studies in other communities.

## Acknowledgements

This study was supported by Agencia Nacional de Promoción Científica y Tecnológica (PICT 2012-2095 PI M. Marchese, and PICT 2016-0465 PI M. Devercelli), Consejo Nacional de Investigaciones Científicas y Técnicas (CONICET). H. Sarmento work was supported by Conselho Nacional de Desenvolvimento e Pesquisa Tecnológica (CNPq process 309514/2017-7) and Fundação de Amparo à Pesquisa do Estado de São Paulo (FAPESP process 2014/14139-3). We thank C. Debonis, E. Creus, and M. Piacenza for their field assistance, and A. Dabin for bioinformatic support. We thank the members of Metacommunity Project particularly M. Marchese, P. Scarabotti, D. Borzone, M. Licursi, for the long discussions that largely improved the manuscript.

## Author contributions

PH and MD conceived and designed the study; MD conducted the samplings with the help of GM; PH and SM performed molecular analyzes; GM performed chemical analyses; PH ran statistical analyses; PH and MD wrote the manuscript with substantial contributions from FU and HS; all authors edited the manuscript.

The authors declare that they have no conflict of interest.

Supplementary information is available.

