## Supplementary Information for "Environmental heterogeneity ruling the action of ecological processes over the bacterial metacommunity assembly"

### Supplementary methods

#### *Environmental data collection and analysis*

Subsurface water samples were collected by duplicate at the center of the lotic environments and in the pelagic zone of the lakes. Depth, pH, conductivity, temperature, and dissolved oxygen (HANNA checkers) were measured at sampling sites.

Water samples were transported on ice and in darkness to the laboratory. Turbidity (formazin turbidity units, FTU) was measured with a HACH DR2000 spectrophotometer at 450 nm wavelength. A variable volume of water (200–1,000 mL) was filtered through Whatman GF/C glass fibre filters, which were stored at −20 °C up to 3 weeks. Chlorophyll-*a* (a proxy of phytoplankton biomass) was extracted from the filters with acetone (90%) and spectrophotometrically estimated according to Lorenzen's method [1]. Filtered water samples were passed through Millipore filters (pore size: 0.45 µm) for colorimetric determination of dissolved components. Nitrate ($NO_3^-$) was determined by reduction of nitrate with hydrazine sulfate and subsequent determination of nitrite by diazotizing with sulfanilamide and coupling with N-(1-naphthyl)-ethylenediamine dihydrochloride [2], ammonium ($NH_4^+$) by the indophenol blue method, soluble reactive phosphorus (SRP) by the ascorbic acid method, and dissolved silica (DSi) by the molybdosilicate method.

Chromophoric dissolved organic matter (CDOM) was analyzed using a HACH DR5000 ultraviolet–visible spectrophotometer and 1-cm quartz cuvettes. Absorption spectra (200–700 nm) were measured with 1-nm resolution. Filtered Milli-Q water was used as a baseline. As the absorbance of the CDOM was assumed to be equal to zero above 700 nm, the absorbance at this wavelength was subtracted from all the rest to correct offsets [3]. Absorption coefficients ($m^{-1}$) at each wavelength were calculated according to Kirk [4]:

$$a_\lambda = 2.303 \times A_\lambda \div l$$

where $a_\lambda$ is the CDOM absorption coefficient at wavelength $\lambda$, $A_\lambda$ is the corrected absorbance at wavelength $\lambda$, and $l$ is the cuvette path length in m. $a_{440}$ was used as a measure of CDOM concentration [4]. In addition, the spectral slope for the interval of 275–295 nm ($S_{275\text{-}295}$) was calculated using linear regression of Ln-transformed $a$ spectra. It has been shown to inversely covary with CDOM molecular weight. Therefore, it was used to characterize shifts in CDOM quality and sources [5].

*DNA extraction*

Genomic DNA from filters was extracted using a CTAB protocol [6]. Briefly, warm CTAB lysis buffer was added to the filters and incubated at 60 °C for 30 min. Two purification steps were performed adding 0.7 ml of chloroform-isoamilic alcohol (24:1) and centrifuging at 14 000 rpm for 10 min. Subsequently, the DNA was precipitated in cold isopropanol and centrifuged at 14 000 rpm for 30 min. Finally, a washing step of the pellet using cold ethanol (80%) was performed, and the extracted DNA was air-dried and re-suspended in 40 µl of TE buffer.

*Sequence Data analysis*

The reads were first analyzed for error correction using the algorithms based on Hamming graphs and Bayesian subclustering (BAYES HAMMER tool) [7] implemented in SPAdes v3.5.0 [8]. Then, the forward and reverse sequences were assembled using the function fastq_mergepairs from USEARCH-v10 [9]. The minimum overlap length was set to 20 bp and assemblage sequences with less than 100 nucleotides were discarded; the rest of the parameters were used as the default. The reads' quality was determined using fastq_filter in USEARCH-v10 [9] using maxee value of 0.5 and a minimum length of 100 pb. From reads that passed the quality control, singletons were removed. The taxonomic assignation of Operational taxonomic units (zOTUs) was done by BLAST [10], using the SILVA database (SSU Ref 132 NR 99) as a reference. Finally, the zOTU table was constructed using the otutab function in USEARCH-v10 [9].

*Estimation of zOTU's niche and phylogenetic distances*

The zOTU's niche distances (i.e., differential environment optima requirements of each zOTUs) were estimated for each hydrological phase with respect to nine environmental variables: DO, turbidity, conductivity, pH, SRP, $NO_3^-$, $NH_4^+$, CDOM concentration and CDOM molecular weight. We calculated the zOTUs relative-abundance–weighted mean value for each environmental variable as following: for each variable in each sample we registered all records of a given zOTU and multiply its abundance by the variable's value; these values were summed across all samples where a zOTU was found and divided by the zOTU total abundance value. Then we generated a matrix containing all estimating environmental optima, normalized as standard normal deviates, and calculated the distance matrix using Euclidean index [11, 12].

The zOTU phylogenetic distances were computed based on maximum likelihood trees constructed for each hydrological phase (Ape package in the R environment [13]). The sequences were aligned using MAFFT v.7 [14], and the phylogenetic trees were constructed using the Randomized Axelerated Maximum Likelihood (RAxML) v.8 program [15] with the GTR-GAMMA model considering 500 bootstraps.

The Mantel correlogram analyses were run with 999 permutations (Vegan v.2.0.9 package [16] in R), using 50 phylogenetic distance bins, and a progressive Bonferroni correction.

**Supplementary tables**

**Table S1:** Mean values and variation ranges of environmental variables measured at the Paraná fluvial systems in four hydrological phases and at different type of environments.

and variation ranges of environmental variables measured at the Paraná fluvial systems in four hydrological phases and at different type of environments.

| *Environmental Type*[a] | *temperatura* °C | *conductivity* $\mu S\ cm^{-1}$ | *dissolved oxygen* $mg\ l^{-1}$ | *pH* | *Secchi* m | *turbidity* FTU | *SPR*[b] $\mu g\ l^{-1}$ | *nitrate* $\mu g\ l^{-1}$ | *ammonium* $\mu g\ l^{-1}$ | $A_{440}$[c] $m^{-1}$ | $S_{275\text{-}295}$[d] slope | *chlorophyll-* $\mu g\ l^{-1}$ |
|---|---|---|---|---|---|---|---|---|---|---|---|---|
| **IL** | 26.4 (22.4-28.9) | 130 (81-242) | 6.8 (0.2-12.3) | 7.8 (6.3-9.6) | 0.49 (0.12-1.1) | 26 (4-65) | 50 (13-187) | 67 (0-169) | 46 (5-155) | 2.8 (1.6-5.8) | 0.0156 (0.0153-0.0160) | 14 (3.2-47) |
| **CL** | 26.6 (25.5-27.8) | 82 (80-83.4) | 3.5 (1-5.2) | 7.5 (7.4-7.6) | 0.62 (0.31-1.1) | 8 (5-11) | 15 (6-22) | 121 (81-181) | 70 (53-92) | 1.8 (1.6-1.9) | 0.0154 (0.0152-0.0157) | 6.2 (4.9-8) |
| **SC** | 27.6 (27.5-27.8) | 131 (78-184) | 6.1 (5.1-7) | 7.3 (7.3-7.4) | 0.31 (0.3-0.32) | 31 (14-51) | 47 (35-59) | 317 (203-432) | 75 (30-119) | 2.2 (1.4-3) | 0.0173 (0.0138-0.0201) | 6.8 (4.7-8.8) |
| **MC** | 29.6 (24.5-36.2) | 146 (72-286) | 7 (6.5-7.4) | 7.4 (7-7.8) | 0.32 (0.29-0.36) | 26 (12-54) | 50 (14-111) | 371 (206-459) | 51 (27-68) | 2.7 (1.6-3.9) | 0.0158 (0.0153-0.0163) | 3.6 (3-4.8) |
| **IL** | 22.7 (21.1-25.5) | 109 (56-319) | 7.1 (1-15) | 7.2 (6.3-8.8) | 0.57 (0.16-1.1) | 32 (3-67) | 9 (0-27) | 34 (0-105) | 16 (2-41) | 2.4 (2-3.4) | 0.0157 (0.0148-0.0163) | 26 (4.4-120.5 |
| **CL** | 23.3 (22.7-24.2) | 107 (96-117) | 5.1 (5-5.2) | 7 (6.9-7.1) | 0.72 (0.14-1.24) | 35 (11-77) | 1 (0-2) | 32 (1-95) | 16 (3-44) | 1.5 (1.1-1.8) | 0.0150 (0.0139-0.0158) | 5.3 (4.4-6) |
| **SC** | 23.7 (22.9-24.7) | 99 (75-130) | 7.4 (5.8-8.2) | 7 (6.9-7.2) | 0.16 (0.11-0.19) | 91 (76-100) | 15 (8-24) | 234 (108-310) | 36 (24-69) | 2.1 (1.5-3.1) | 0.0182 (0.0158-0.0212) | 2.6 (1.8-3.2) |
| **MC** | 23.6 (23.2-24.1) | 179 (77-377) | 8.1 (6.7-9.8) | 7.4 (7.3-7.5) | 0.12 (0.1-0.16) | 146 (86-178) | 40 (22-58) | 327 (300-361) | 21 (2-37) | 2.1 (1.4-3.2) | 0.0174 (0.0170-0.0178) | 2.1 (0.8-4.5) |
| **IL** | 17.1 (15-18) | 85 (69-127) | 6.1 (1.9-11.3) | 6.3 (5.8-6.9) | 0.59 (0.11-1.09) | 15 (9-21) | 58 (40-103) | 43 (0-74) | 14 (2-23) | 4.8 (3.8-6.8) | 0.0130 (0.0129-0.0133) | 31.9 (4.1-87.6) |
| **CL** | 17.9 (17.5-18.5) | 69 (68-70) | 7.7 (5.9-9.3) | 6.6 (6-7) | 0.93 (0.6-1.28) | 10 (6-14) | 40 (36-43) | 69 (25-159) | 11 (2-25) | 4 (3.7-4.2) | 0.0132 (0.0128-0.0140) | 2.7 (2.2-3.3) |
| **SC** | 19 (19-19) | 70 (69-70) | 6.9 (6.0-7.7) | 6.3 (6.2-6.3) | 0.55 (0.45-0.65) | 19 (19-19) | 30 (26-35) | 97 (97-98) | 18 (16-19) | 3.9 (3.4-4.2) | 0.0135 (0.0133-0.0137) | 2.6 (2.5-2.6) |
| **MC** | 19.5 (19-20) | 67 (66-69) | 10.2 (9-11.3) | 7 (6.9-7) | 0.33 (0.33-0.33) | 32 (31-32) | 30 (17-44) | 279 (231-328) | 29 (22-36) | 4.3 (3.7-4.8) | 0.0133 (0.0130-0.0136) | 1.7 (1.4-1.9) |
| **IL** | 28.3 (26.8-30) | 65 (61-70) | 3.1 (1.7-5.3) | 6.6 (6.2-7.2) | 1.05 (0.55-1.4) | 11 (7-17) | 35 (31-41) | 177 (112-251) | 19 (7-35) | 3.4 (2.6-4.3) | 0.0128 (0.0127-0.0130) | 2.5 (1.4-3.2) |
| **CL** | 28.3 (26.8-30) | 65 (61-70) | 3.1 (1.7-5.3) | 6.6 (6.2-7.2) | 1.05 (0.55-1.4) | 11 (7-17) | 35 (31-41) | 177 (112-251) | 19 (7-35) | 3.4 (2.6-4.3) | 0.0128 (0.0128-0.0129) | 2.5 (1.4-3.2) |
| **SC** | 29.5 (29-30) | 64 (61-66) | 3 (2.5-3.4) | 6.4 (6.1-6.7) | 1.09 (1.05-1.13) | 11 (11-12) | 29 (28-30) | 158 (123-193) | 16 (7-25) | 3 (2.5-3.4) | 0.0125 (0.0112-0.0133) | 3.1 (2.2-4) |
| **MC** | 26.5 (25-28) | 65 (65-66) | 5.2 (4.7-5.8) | 6.3 (6.3-6.4) | 0.62 (0.46-0.78) | 21 (13-29) | 32.88 (24.57-41.19) | 297 (293-301) | 29 (19-38) | 4 (4-4) | 0.0130 (0.0127-0.0132) | 1.6 (1.4-1.8) |

ected lakes, SC=secondary channels, MC=main channels. [b]soluble reactive phosphorous. [c] Absorption coefficients of chromophoric dissolved organic matter at 440 nm wavelength. [d] spectral slope for
anic matter (275-295 nm wavelength).

**Table S2:** PERMANOVA test performed to determine whether pair-wise environmental conditions and baterial structure (zOTUs abundance) differed among hydrological phases at the Paraná fluvial system. We used Euclidean similarity index based on standardized abiotic variables data with 9 999 permutations, and pairwise tests were then used to compare differences between hydrological phases with the Bonferroni *P*-values correction method (bold letters).

| | **Abiotic variables** | **Bacteria structure** |
|---|---|---|
| Total SS | 522 | |
| Within-group SS | 358 | |
| Pseudo-F | 8.4 | |
| P-values | **0.001** | **0.05** |
| pLW vs LW | **0.0216** | 0.9786 |
| pLW vs HW | **0.0006** | **0.0006** |
| pLW vs eHW | **0.0006** | **0.0006** |
| pLWs vs HW | **0.0006** | **0.0012** |
| LW vs eHW | **0.0006** | **0.0006** |
| HW vs eHW | **0.0006** | **0.0006** |

### Supplementary figures

**Fig. S1**: Mantel correlograms (Pearson correlations) between zOTU niche distances and phylogenetic distances for each hydrological phase of the Paraná fluvial system, with 9 999 permutations. Significant correlations (P<0.05, solid circles) were detected over short phylogenetic distances indicating that closely related taxa are more similar in their habitat preferences than the distantly related taxa. For each phylogenetic distance, bin phylogenetic distances were normalized to vary between 0 and 1 before analysis.

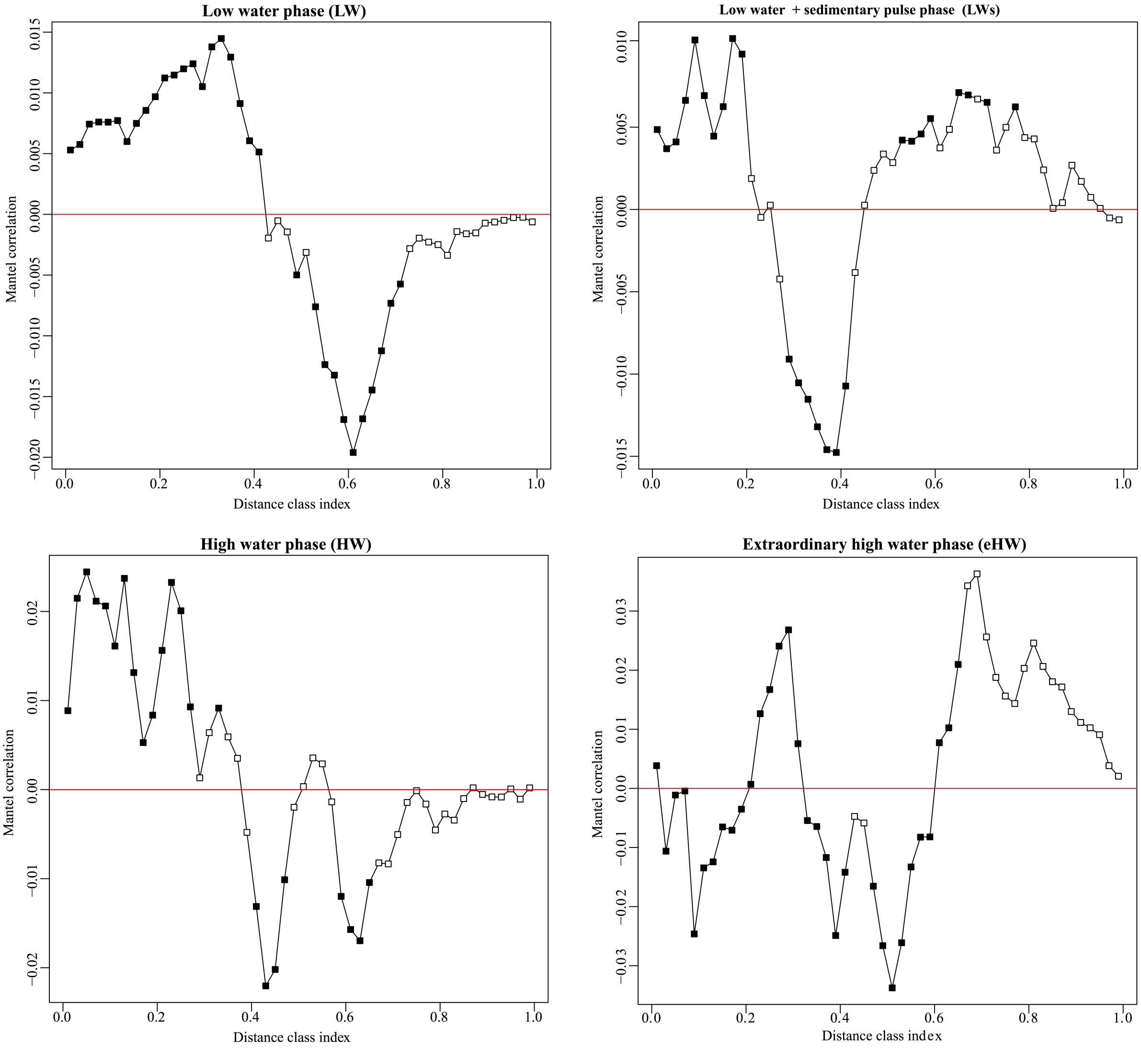

**Fig. S2**: Rarefaction curve for the 4 shallowly sequenced bacterial communities of the Paraná fluvial system in the four shallowly sequenced samples (Vegan package in R). The zOTU accumulation curves did not reach the plateau, indicating that the sequencing effort captured a large fraction, but not all bacterial richness. The different lengths of the curves reflect the variable sequencing coverage per sample.

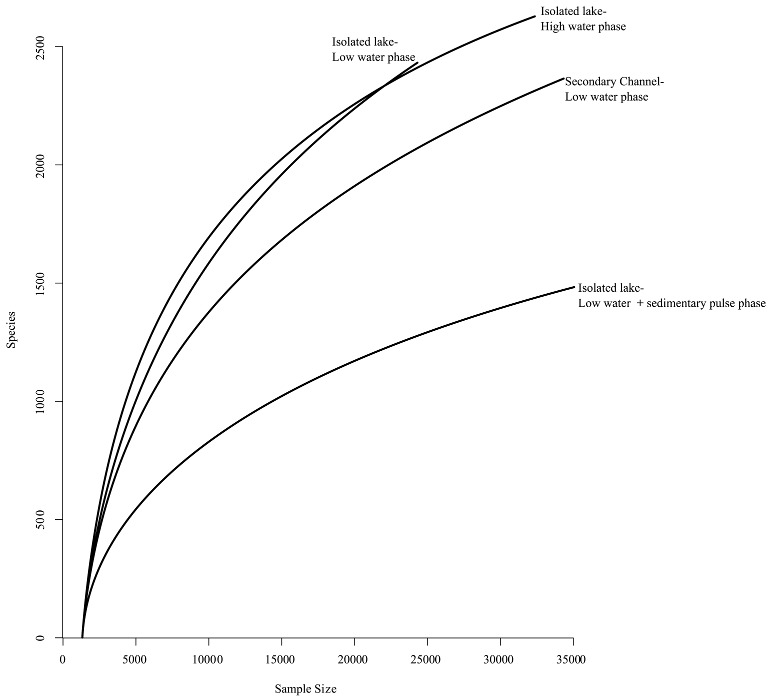

**Fig. S3**: Rank curve of bacterial metacommunities of the Paraná fluvial system in each hydrological pase (Vegan package in R) showing a typical pattern where the abundant fraction was represented by few zOTUs (22 to 25), accounting for more than 80% of total reads.

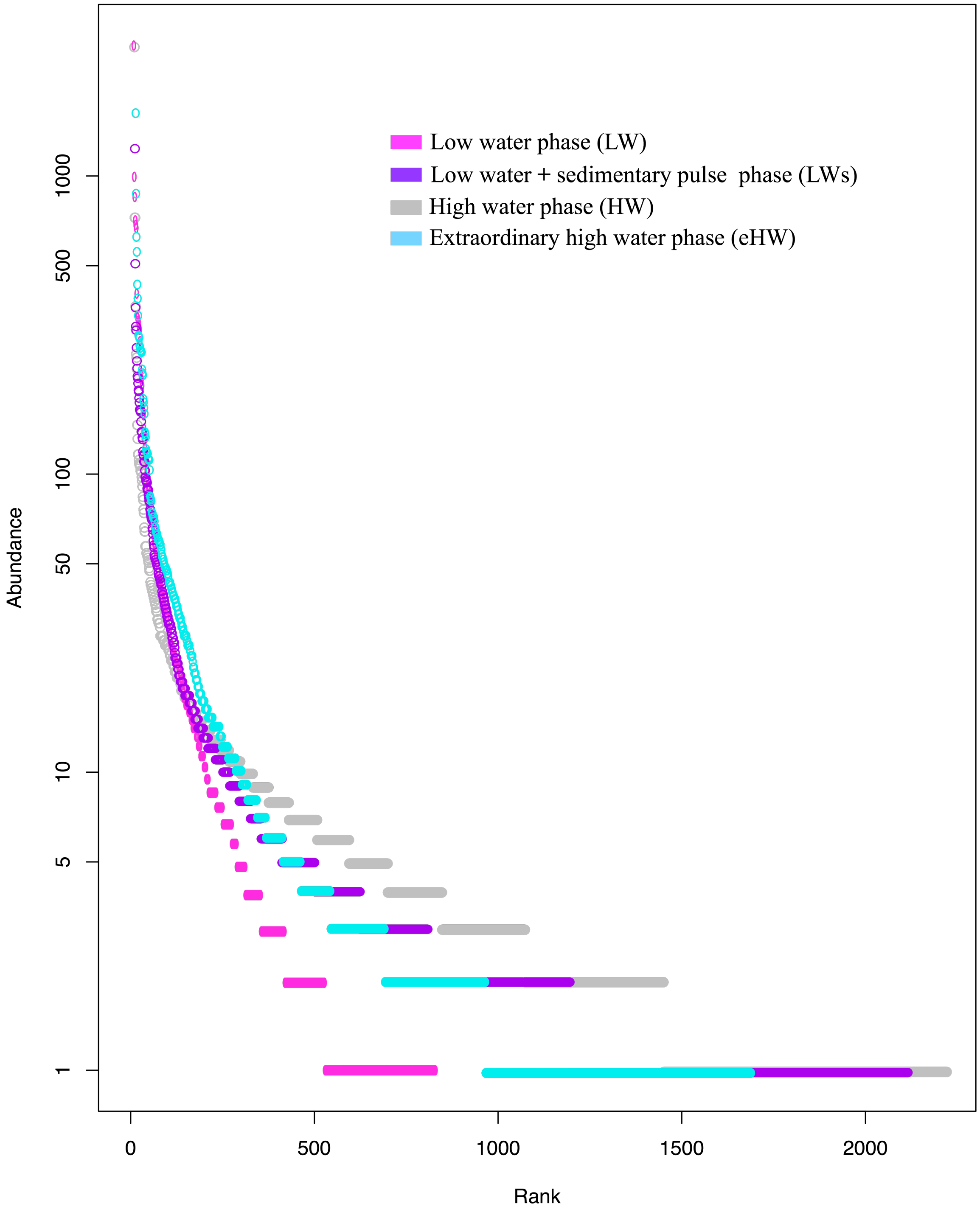

**Fig. S4:** Relative abundance of bacterial taxonomic groups at each hydrological phase of the Paraná fluvial system. LW: low water, LWs: low water with sedimentological pulse, HW: high water, eHW: extraordinary high water.

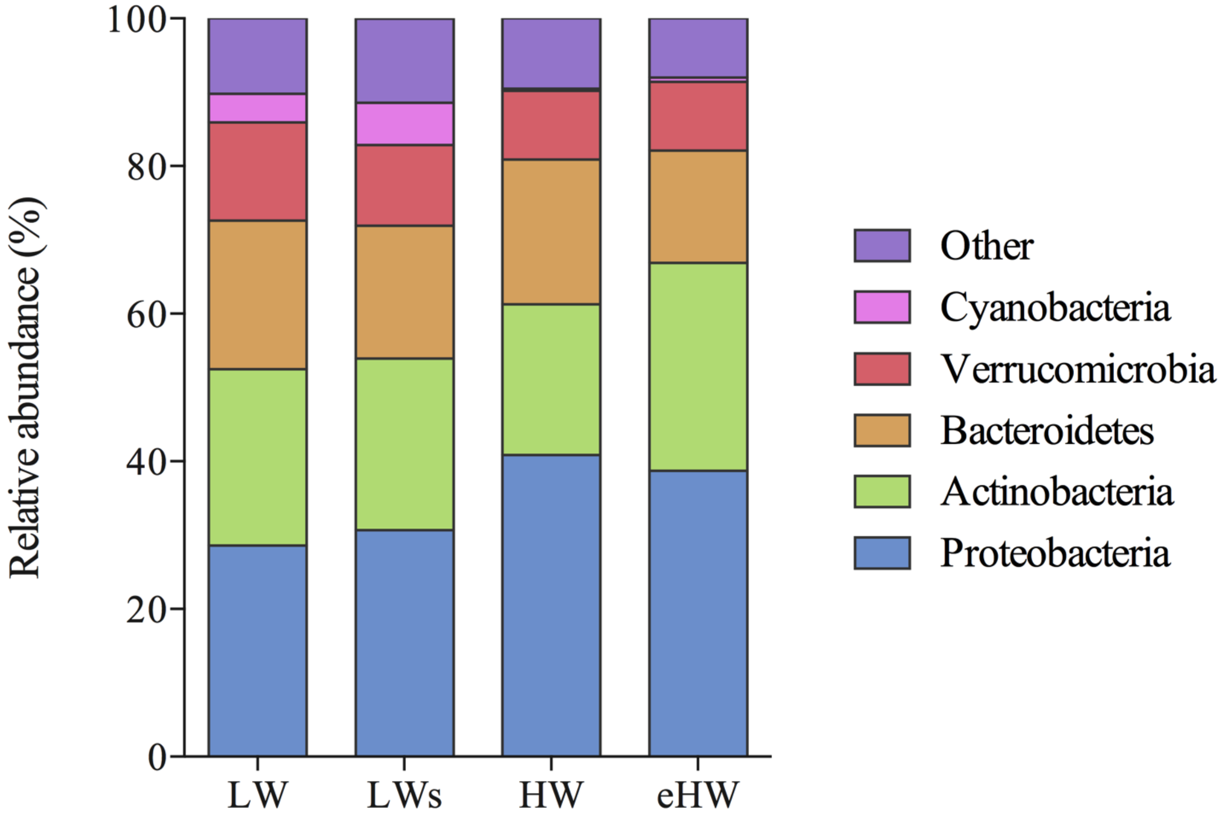

**Fig. S5:** Number of keystone taxa per taxonomic group at each hydrological phase of the Paraná fluvial system. LW: low water, LWs: low water with sedimentological pulse, HW: high water, eHW: extraordinary high water.

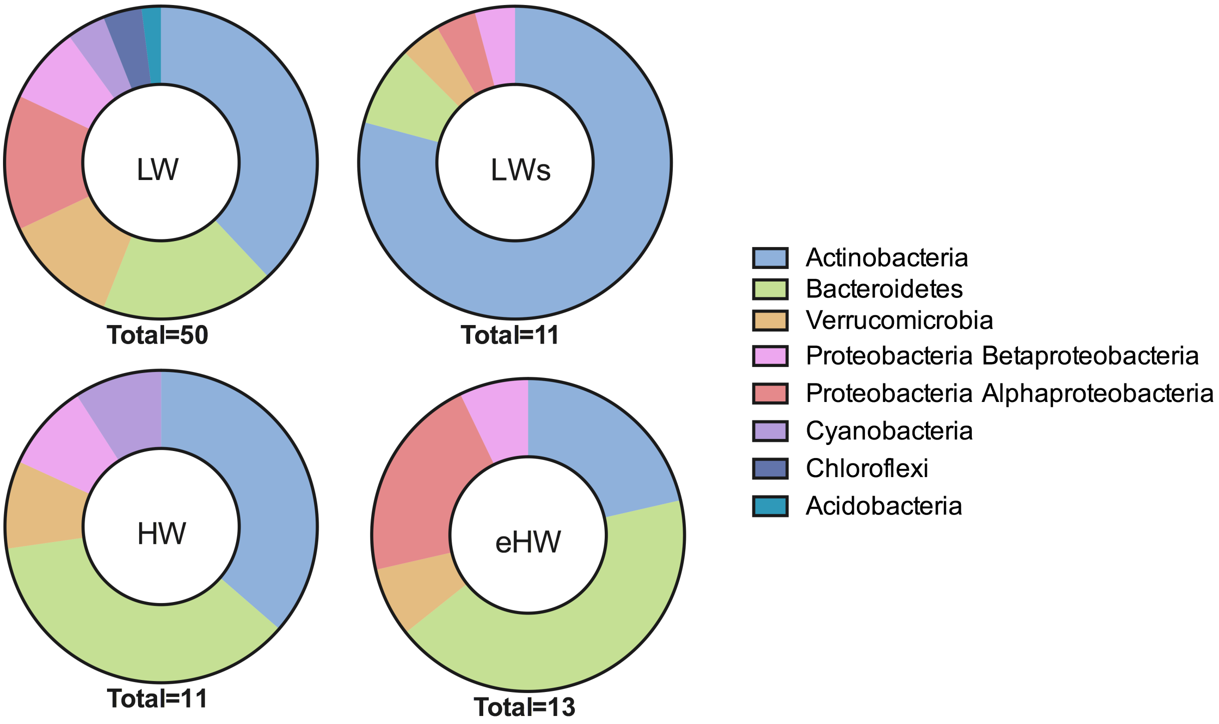

8. Nurk S, Bankevich A, Antipov D, Gurevich AA, Korobeynikov A, Lapidus A, et al. Assembling single-cell genomes and mini-metagenomes from chimeric MDA products.

*J Comput Biol* 2013; **20**: 714–737.

9. Edgar RC, Flyvbjerg H. Error filtering, pair assembly and error correction for next-generation sequencing reads. *Bioinformatics* 2015; **31**: 3476–3482.

10. Altschul SF, Gish W, Miller W, Myers EW, Lipman DJ. Basic local alignment search tool. *J Mol Biol* 1990; **215**: 403–410.

11. Stegen JC, Lin X, Fredrickson JK, Chen X, Kennedy DW, Murray CJ, et al. Quantifying community assembly processes and identifying features that impose them. *ISME J* 2013; **7**: 2069–2079.

12. Dini-Andreote F, Stegen JC, Van Elsas JD, Salles JF. Disentangling mechanisms that mediate the balance between stochastic and deterministic processes in microbial succession. *Proc Natl Acad Sci U S A* 2015; **112**: E1326–E1332.

13. R Development Core Team 3.0.1. A Language and Environment for Statistical Computing. *R Found Stat Comput* . 2013.

14. Katoh K, Standley DM. MAFFT Multiple Sequence Alignment Software Version 7: Improvements in Performance and Usability. *Mol Biol Evol* 2013; **30**: 772–780.

15. Stamatakis A. RAxML version 8: A tool for phylogenetic analysis and post-analysis of large phylogenies. *Bioinformatics* 2014; **30**: 1312–1313.

16. Oksanen J, Blanchet FG, Kindt R, Legendre P, Minchin PR, O'Hara RB, et al. Vegan: Community Ecology Package. R package version 2.0-9. 2013.
